# mRNA-1273 and BNT162b2 mRNA vaccines have reduced neutralizing activity against the SARS-CoV-2 Omicron variant

**DOI:** 10.1101/2021.12.20.473557

**Authors:** Venkata-Viswanadh Edara, Kelly E. Manning, Madison Ellis, Lilin Lai, Kathryn M. Moore, Stephanie L. Foster, Katharine Floyd, Meredith E. Davis-Gardner, Grace Mantus, Lindsay E. Nyhoff, Sarah Bechnak, Ghina Alaaeddine, Amal Naji, Hady Samaha, Matthew Lee, Laurel Bristow, Laila Hussaini, Caroline Rose Ciric, Phuong-Vi Nguyen, Matthew Gagne, Jesmine Roberts-Torres, Amy R. Henry, Sucheta Godbole, Arash Grakoui, Marybeth Sexton, Anne Piantadosi, Jesse J. Waggoner, Daniel C. Douek, Evan J. Anderson, Nadine Rouphael, Jens Wrammert, Mehul S. Suthar

## Abstract

The BNT162b2 (Pfizer-BioNTech) and mRNA-1273 (Moderna) vaccines generate potent neutralizing antibodies against severe acute respiratory syndrome coronavirus 2 (SARS-CoV-2). However, the global emergence of SARS-CoV-2 variants with mutations in the spike protein, the principal antigenic target of these vaccines, has raised concerns over the neutralizing activity of vaccine-induced antibody responses. The Omicron variant, which emerged in November 2021, consists of over 30 mutations within the spike protein. Here, we used an authentic live virus neutralization assay to examine the neutralizing activity of the SARS-CoV-2 Omicron variant against mRNA vaccine-induced antibody responses. Following the 2nd dose, we observed a 30-fold reduction in neutralizing activity against the omicron variant. Through six months after the 2nd dose, none of the sera from naïve vaccinated subjects showed neutralizing activity against the Omicron variant. In contrast, recovered vaccinated individuals showed a 22-fold reduction with more than half of the subjects retaining neutralizing antibody responses. Following a booster shot (3rd dose), we observed a 14-fold reduction in neutralizing activity against the omicron variant and over 90% of boosted subjects showed neutralizing activity against the omicron variant. These findings show that a 3rd dose is required to provide robust neutralizing antibody responses against the Omicron variant.

## Main

The BNT162b2 (Pfizer-BioNTech) and mRNA-1273 (Moderna) vaccines generate potent and durable neutralizing antibody responses against the severe acute respiratory syndrome coronavirus 2 (SARS-CoV-2)^1–3^. The global emergence of SARS-CoV-2 variants with mutations in the spike protein, the principal antigenic target of these vaccines, has raised concern regarding the effectiveness of these vaccines. We previously found that mRNA vaccine-induced antibody responses have reduced neutralizing activity against the B.1.351 (Beta), and to a lesser extent, B.1.617.2 (Delta) variants^4–7^.

The B.1.1.529 (Omicron) variant emerged in November 2021 and has rapidly spread throughout the world. We isolated the B.1.1.529 variant from a residual mid turbinate swab collected from a returning traveler from South Africa (hCoV-19/USA/GA-EHC-2811C/2021). Relative to the WA1/2020 virus (nCoV/USA_WA1/2020; spike 614D), the B.1.1.529 variant contains several mutations within the spike protein (A67V, Δ69-70, T95I, G142D, Δ143-145, Δ212, N211I, +214EPE, G339D, S371L, S373P, S375F, K417N, N440K, G446S, S477N, T478K, E484A, Q493R, G496S, Q498R, N501Y, Y505H, T547K, D614G, H655Y, N679K, P681H, N764K, D796Y, N856K, Q954H, N969K, L981F). As a comparator, we included the B.1.351 (Beta) variant in our neutralization assay. The Beta variant has mutations within the spike protein at amino acid residues L18F, D80A, D215G, L241-, L242-, A243-, K417N, E484K, N501Y, D614G, A701V. We used an *in vitro* authentic live virus Focus Reduction Neutralization Test (FRNT)^8^ on Vero-TMPRSS2 cells to perform a cross-sectional analysis of neutralizing antibody response in serum (**Supplementary Tables 1-4**) from three naïve mRNA vaccinated cohorts, which includes 2-4 weeks after the primary series (n=24; n=11 Moderna; n=13 Pfizer-BioNTech), 6 months after the primary series (n=25; n=8 Moderna; n=17 Pfizer-BioNTech), and 1-4 weeks after a single booster dose (n=52; n=17 Moderna 3rd dose; n=35 Pfizer-BioNTech 3rd dose), and a COVID-19 recovered then mRNA vaccinated cohort (6 months post-2nd dose; n=37; n=13 Moderna; n=24 Pfizer-BioNTech).

In Moderna or Pfizer-BioNTech vaccinated individuals, we found profoundly reduced neutralizing activity against the B.1.1.529 variant as compared to either the WA1/2020 strain or B.1.351 variant. Following primary mRNA vaccination in naïve individuals, the FRNT Geometric Mean Titers (GMTs) were 520 for WA1, 97 for B.1.351 and 17 for B.1.1.529 and corresponded to a 5.4-fold and 30-fold reduction as compared to WA1, respectively (**Fig. 1a**). Further, only 21% of the subjects showed neutralizing antibody titers against the B.1.1.529 variant.

**Figure 1.**
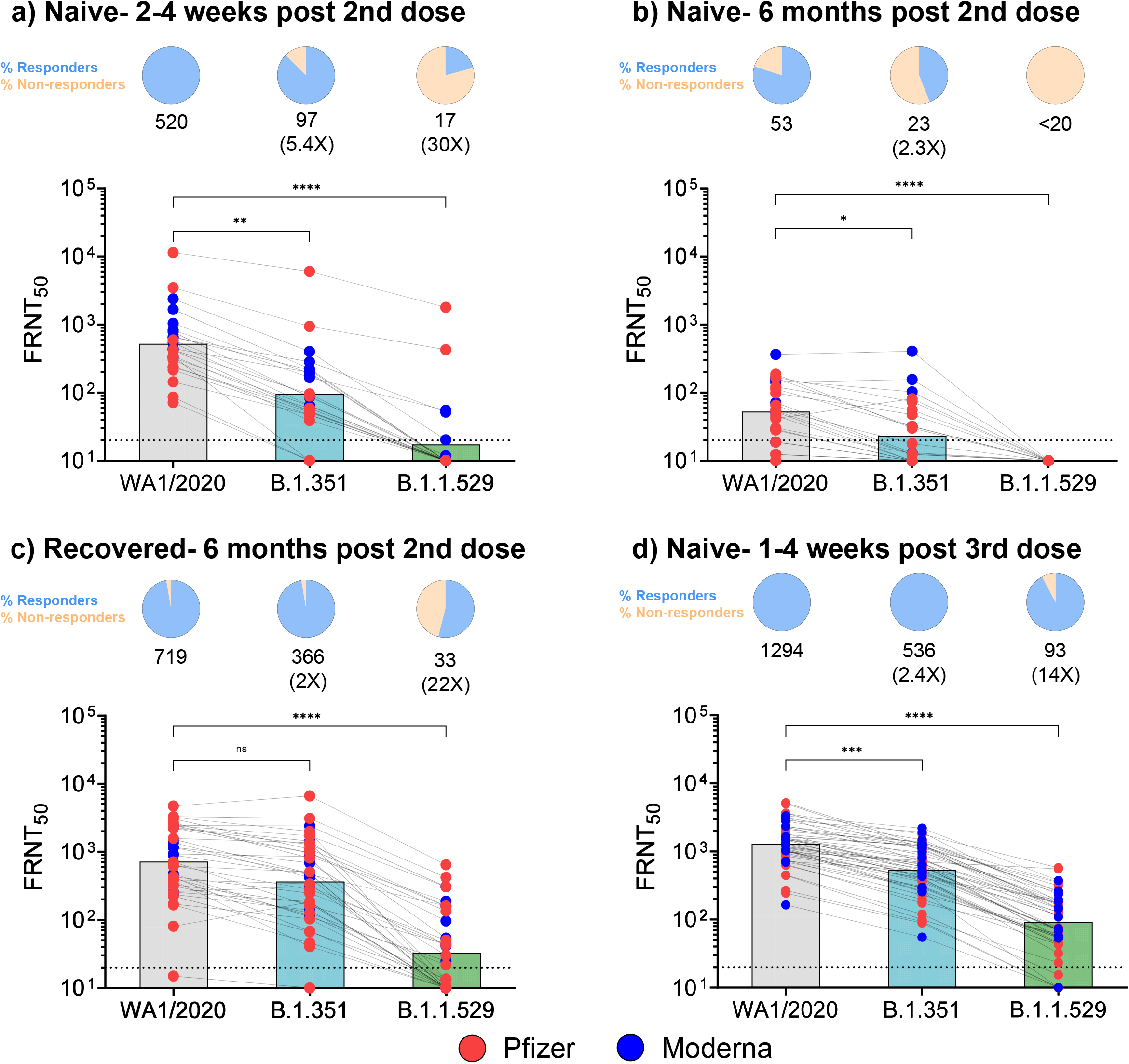
Neutralization antibody responses against WA1/2020, B.1.351, and B.1.1.529 SARS-CoV-2 variants post-mRNA vaccination. Data from the following cohorts are shown: **A)** Naïve individuals 2-4 weeks post-2nd dose (n=24); **B)** Naïve individuals 6-months post-2nd dose (n=25); **C)** Recovered individuals that received the primary mRNA vaccination series (n=37); and **D)** Naïve vaccinated individuals that received a 3rd dose (n=52). In panels A-D, the FRNT_50_ GMTs for WA1/2020, B.1.351 and B.1.1.529 are shown with respective fold changes in comparison to the WA1/2020. A pie-chart above each graph shows the frequency of individuals that have titers above (Responders) or below (Non-responders) the limit of detection (LOD). The connecting lines between the variants represents matched serum samples. The horizontal dashed lines along the X-axis indicate the limit of detection (FRNT_50_ GMT= 10). Blue circles represent individuals that received the Moderna mRNA-1273 vaccine as the primary vaccine series and the red circles represent individuals that received the Pfizer-BioNTech BNT162b2 vaccine as the primary vaccine series. Non-parametric were performed using a Kruskal-Wallis test with Dunn’s multiple comparisons test. *p<0.05; **p<0.01; ***p<0.001; ****p<0.0001

Next, we examined the durability of neutralizing antibody responses in vaccinated subjects 6 months after receiving the 2nd dose. This cohort was divided between individuals with no known prior COVID-19 exposure and those that had recovered from COVID-19 and then received the Moderna or Pfizer-BioNTech vaccine. In naïve vaccinated subjects, the GMTs at 6 months were 53 for WA1, 23 for B.1.351, and <20 for B.1.1.529 (**Fig. 1b**). This corresponded to a 2.3-fold reduction against B.1.351 as compared to WA1, however, none of these subjects showed any detectable neutralizing activity against the B.1.1.529 variant. In recovered individuals that received the vaccine, the GMTs were 719 for WA1, 366 for B.1.351 and 33 for B.1.1.529 and corresponded to a 2- and 22-fold reduction as compared to WA1, respectively (**Fig. 1c**). In contrast to naïve vaccinated subjects, 55% of recovered vaccinated subjects retained neutralizing activity against the B.1.1.529 variant 6 months after the 2nd dose.

Next, we examined the impact of the B.1.1.529 variant on neutralizing antibody responses following a single booster dose (3rd dose). Most subjects received the 3rd dose approximately 8 months (median 268 days) after the 2nd dose. Roughly 85% of these subjects received a homologous 3rd dose (primary vaccine vs booster dose) and a few subjects received 100 mcg of mRNA-1273 as the 3rd dose instead of the 50 mcg currently approved as a booster dose. Sampling occurred 1-4 weeks after the booster dose. Across all subjects that received a booster dose (**Fig. 1d**), the GMTs were 1294 for WA1, 536 for B. 1.351, and 93 for B.1.1.529. This corresponded to a reduction of 2.4-fold and 14-fold in neutralizing activity as compared to WA1 and over 90% of subjects retained neutralizing activity against the B.1.1.529 variant.

Our findings show that the B.1.1.529 variant has a significant impact on the neutralizing activity against mRNA vaccine-induced responses. By 6 months, all naïve vaccinated and a majority of recovered vaccinated subjects lost neutralizing activity against the B.1.1.529 variant. However, following a booster dose (3rd dose), a vast majority of subjects retained neutralizing activity against B.1.1.529. These findings support the need for a booster dose to maintain neutralizing activity against the B.1.1.529 variant.

Limitations of this study include: 1) small sample size; 2) selection bias in sampling for the 3rd dose (Pfizer-BioNTech 3rd dose received EUA approval prior to Moderna); 3) this study was a cross-sectional analysis of subjects that received either the primary or booster mRNA vaccines; 4) we are not able to link the clinical outcomes with the neutralization findings; 5) we cannot exclude that some of the naïve individuals were exposed to COVID-19 but were asymptomatic; and 6) this study did not evaluate T cell immunity which likely plays a role in protection against COVID-19.

## Methods

### Viruses and cells

VeroE6-TMPRSS2 cells were generated and cultured as previously described^7^. nCoV/USA_WA1/2020 (WA/1), closely resembling the original Wuhan strain and resembles the spike used in the mRNA-1273 and Pfizer-BioNTech vaccine, was propagated from an infectious SARS-CoV-2 clone as previously described^9^. icSARS-CoV-2 was passaged once to generate a working stock. The B.1.351 variant isolate, kindly provided by Dr. Andy Pekosz (John Hopkins University, Baltimore, MD), was propagated once in VeroE6-TMPRSS2 cells to generate a working stock. hCoV19/EHC_C19_2811C (herein referred to as the B.1.1.529 variant) was derived from a mid-turbinate nasal swab collected in December 2021. This SARS-CoV-2 genome is available under GISAID accession number EPI_ISL_7171744. Using VeroE6-TMPRSS cells, the B.1.1.529 variant was plaque purified directly from the nasal swab, propagated once in a 12-well plate, and expanded in a confluent T175 flask to generate a working stock. All viruses used in this study were deep sequenced and confirmed as previously described^7^.

### Samples

At Emory University, collection and processing were performed under the University Institutional Review Board protocols #00045821, #00002061 and #00022371 at the Emory Hope Clinic and Emory Children’s Center. Naïve and non-naïve adults ≥18 years were enrolled if they met eligibility criteria for these umbrella protocols and provided informed consent. Convalescent samples were a convenience sample of individuals with confirmed mild or moderate COVID-19 (March-August 2020)^6^. These participants subsequently received vaccination with 2 doses of Pfizer BNT162b2 or Moderna mRNA1273 and their sera or plasma samples were collected 6 months after vaccination. Three other cohorts of naïve participants were enrolled after receiving either mRNA vaccines and their sera or plasma were collected at the following timepoints: 1) 2-4 weeks after primary series; 2) 6 months after primary series and 3) 1-4 weeks after single dose boost (85% were homologous boosts).

### Focus Reduction Neutralization Test

FRNT assays were performed as previously described^6–8^. Briefly, samples were diluted at 3-fold in 8 serial dilutions using DMEM (VWR, #45000-304) in duplicates with an initial dilution of 1:10 in a total volume of 60 μl. Serially diluted samples were incubated with an equal volume of WA1/2020, B.1.351, or B.1.1.529 (100-200 foci per well based on the target cell) at 37° C for 45 minutes in a round-bottomed 96-well culture plate. The antibody-virus mixture was then added to VeroE6-TMPRSS2 cells and incubated at 37°C for 1 hour. Post-incubation, the antibody-virus mixture was removed and 100 μl of pre-warmed 0.85% methylcellulose (Sigma-Aldrich, #M0512-250G) overlay was added to each well. Plates were incubated at 37° C for 18 hours and the methylcellulose overlay was removed and washed six times with PBS. Cells were fixed with 2% paraformaldehyde in PBS for 30 minutes. Following fixation, plates were washed twice with PBS and permeabilization buffer (0.1% BSA [VWR, #0332], Saponin [Sigma, 47036-250G-F] in PBS) was added to permeabilized cells for at least 20 minutes. Cells were incubated with an anti-SARS-CoV spike primary antibody directly conjugated to Alexaflour-647 (CR3022-AF647) for up to 4 hours at room temperature. Cells were washed three times in PBS and foci were visualized on Cytation7.

### Quantification and Statistical Analysis

Antibody neutralization was quantified by counting the number of foci for each sample using the Viridot program^10^. The neutralization titers were calculated as follows: 1 - (ratio of the mean number of foci in the presence of sera and foci at the highest dilution of respective sera sample). Each specimen was tested in duplicate. The FRNT-50 titers were interpolated using a 4-parameter nonlinear regression in GraphPad Prism 9.2.0. Samples that do not neutralize at the limit of detection at 50% are plotted at 10 and was used for geometric mean and fold-change calculations.

## References

1. El Sahly, H.M., et al. Efficacy of the mRNA-1273 SARS-CoV-2 Vaccine at Completion of Blinded Phase. N Engl J Med (2021).

2. Baden, L.R., et al. Efficacy and Safety of the mRNA-1273 SARS-CoV-2 Vaccine. N Engl J Med 384, 403–416 (2021).

3. Jackson, L.A., et al. An mRNA Vaccine against SARS-CoV-2 - Preliminary Report. N Engl J Med (2020).

4. Pegu, A., et al. Durability of mRNA-1273 vaccine-induced antibodies against SARS-CoV-2 variants. Science (2021).

5. Suthar, M.S., et al. Durability of immune responses to the BNT162b2 mRNA vaccine. 2021.2009.2030.462488 (2021).

6. Edara, V.V., et al. Infection- and vaccine-induced antibody binding and neutralization of the B.1.351 SARS-CoV-2 variant. Cell Host Microbe 29, 516–521 e513 (2021).

7. Edara, V.V., et al. Infection and Vaccine-Induced Neutralizing-Antibody Responses to the SARS-CoV-2 B.1.617 Variants. N Engl J Med 385, 664–666 (2021).

8. Vanderheiden, A., et al. Development of a Rapid Focus Reduction Neutralization Test Assay for Measuring SARS-CoV-2 Neutralizing Antibodies. Curr Protoc Immunol 131, e116 (2020).

9. Xie, X., et al. An Infectious cDNA Clone of SARS-CoV-2. Cell Host Microbe 27, 841–848 e843 (2020).

10. Katzelnick, L.C., et al. Viridot: An automated virus plaque (immunofocus) counter for the measurement of serological neutralizing responses with application to dengue virus. PLoS Negl Trop Dis 12, e0006862 (2018).

