## Supplementary Table 1 for "mRNA-1273 and BNT162b2 mRNA vaccines have reduced neutralizing activity against the SARS-CoV-2 Omicron variant"

**Supplemental Table 1. FRNT50 results from Naïve Moderna or Pfizer vaccinated samples (2-4 weeks post 2nd dose)**

| S. NO | Age | Sex | Initial vaccine | Days Post Dose 2 | WA1/2020 | B.1.351 | B.1.1.529 |
|---|---|---|---|---|---|---|---|
| 1 | 28 | Female | Moderna | 29 | 667 | 215 | 10 |
| 2 | 32 | Male | Moderna | 27 | 832 | 167 | 55 |
| 3 | 35 | Female | Moderna | 28 | 1043 | 191 | 10 |
| 4 | 44 | Female | Moderna | 28 | 325 | 64 | 10 |
| 5 | 37 | Female | Moderna | 28 | 447 | 198 | 10 |
| 6 | 24 | Female | Moderna | 28 | 644 | 284 | 52 |
| 7 | 25 | Female | Moderna | 32 | 542 | 65 | 10 |
| 8 | 26 | Female | Moderna | 28 | 427 | 53 | 10 |
| 9 | 32 | Male | Moderna | 28 | 1673 | 225 | 10 |
| 10 | 32 | Female | Moderna | 28 | 2389 | 403 | 20 |
| 11 | 28 | Female | Moderna | 34 | 782 | 86 | 12 |
| 12 | 63 | Female | Pfizer | 13 | 342 | 10 | 10 |
| 13 | 62 | Female | Pfizer | 14 | 71 | 10 | 10 |
| 14 | 37 | Male | Pfizer | 13 | 144 | 47 | 10 |
| 15 | 28 | Female | Pfizer | 14 | 346 | 53 | 10 |
| 16 | 60 | Female | Pfizer | 16 | 594 | 89 | 10 |
| 17 | 64 | Female | Pfizer | 16 | 215 | 47 | 10 |
| 18 | 44 | Male | Pfizer | 14 | 304 | 39 | 10 |
| 19 | 25 | Male | Pfizer | 14 | 242 | 51 | 10 |
| 20 | 30 | Female | Pfizer | 14 | 3497 | 942 | 429 |
| 21 | 35 | Female | Pfizer | 14 | 11472 | 6047 | 1801 |
| 22 | 41 | Female | Pfizer | 14 | 86 | 10 | 10 |
| 23 | 23 | Female | Pfizer | 15 | 428 | 96 | 10 |
| 24 | 24 | Male | Pfizer | 14 | 234 | 59 | 10 |
