## Supplementary Table 2 for "mRNA-1273 and BNT162b2 mRNA vaccines have reduced neutralizing activity against the SARS-CoV-2 Omicron variant"

**Supplemental Table 2. FRNT50 results from Naïve Moderna or Pfizer vaccinated samples (6 months post 2nd dose)**

| S.NO | Status | Age | Sex | Vax Brand | Days Post Dose 2 | WA1/2020 | B.1.351 | B.1.1.529 |
|---|---|---|---|---|---|---|---|---|
| 1 | Naïve | 22 | Female | Moderna | 177 | 19 | 10 | 10 |
| 2 | Naïve | 64 | Female | Moderna | 176 | 142 | 156 | 10 |
| 3 | Naïve | 30 | Female | Moderna | 191 | 52 | 10 | 10 |
| 4 | Naïve | 27 | Female | Moderna | 176 | 164 | 32 | 10 |
| 5 | Naïve | 25 | Female | Moderna | 140 | 154 | 104 | 10 |
| 6 | Naïve | 37 | Male | Moderna | 171 | 29 | 10 | 10 |
| 7 | Naïve | 44 | Female | Moderna | 174 | 71 | 13 | 10 |
| 8 | Naïve | 30 | Male | Moderna | 179 | 366 | 408 | 10 |
| 9 | Naïve | 55 | Female | Pfizer | 176 | 113 | 32 | 10 |
| 10 | Naïve | 51 | Female | Pfizer | 175 | 10 | 10 | 10 |
| 11 | Naïve | 24 | Male | Pfizer | 178 | 187 | 30 | 10 |
| 12 | Naïve | 23 | Male | Pfizer | 185 | 168 | 74 | 10 |
| 13 | Naïve | 24 | Female | Pfizer | 175 | 12 | 10 | 10 |
| 14 | Naïve | 41 | Female | Pfizer | 181 | 28 | 10 | 10 |
| 15 | Naïve | 30 | Female | Pfizer | 182 | 56 | 18 | 10 |
| 16 | Naïve | 26 | Male | Pfizer | 176 | 66 | 12 | 10 |
| 17 | Naïve | 29 | Male | Pfizer | 180 | 42 | 82 | 10 |
| 18 | Naïve | 56 | Male | Pfizer | 176 | 99 | 48 | 10 |
| 19 | Naïve | 37 | Male | Pfizer | 176 | 31 | 13 | 10 |
| 20 | Naïve | 45 | Female | Pfizer | 132 | 10 | 10 | 10 |
| 21 | Naïve | 37 | Male | Pfizer | 175 | 19 | 10 | 10 |
| 22 | Naïve | 50 | Female | Pfizer | 179 | 54 | 12 | 10 |
| 23 | Naïve | 34 | Female | Pfizer | 183 | 12 | 10 | 10 |
| 24 | Naïve | 62 | Female | Pfizer | 141 | 122 | 56 | 10 |
| 25 | Naïve | 27 | Female | Pfizer | 175 | 46 | 10 | 10 |
