## Supplementary Table 3 for "mRNA-1273 and BNT162b2 mRNA vaccines have reduced neutralizing activity against the SARS-CoV-2 Omicron variant"

**Supplemental Table 3. FRNT50 results from recovered Moderna or Pfizer vaccinated samples (6 months post 2nd dose)**

| S.NO | Status | Age | Sex | Days PSO | Vax Brand | Days Post Dose 2 | WA1/2020 | B.1.351 | B.1.1.529 |
|---|---|---|---|---|---|---|---|---|---|
| 1 | Recovered | 66 | Female | 296 | Moderna | 179 | 1375 | 419 | 10 |
| 2 | Recovered | 52 | Male | 290 | Moderna | 176 | 1539 | 481 | 10 |
| 3 | Recovered | 38 | Male | 296 | Moderna | 192 | 2288 | 1443 | 54 |
| 4 | Recovered | 69 | Female | 300 | Moderna | 178 | 1154 | 479 | 10 |
| 5 | Recovered | 61 | Male | 285 | Moderna | 175 | 2437 | 974 | 96 |
| 6 | Recovered | 54 | Female | 294 | Moderna | 175 | 935 | 853 | 188 |
| 7 | Recovered | 56 | Male | 354 | Moderna | 178 | 412 | 117 | 13 |
| 8 | Recovered | 64 | Female | 307 | Moderna | 174 | 896 | 466 | 160 |
| 9 | Recovered | 66 | Male | 261 | Moderna | 233 | 262 | 311 | 44 |
| 10 | Recovered | 57 | Male | 334 | Moderna | 177 | 2440 | 2401 | 309 |
| 11 | Recovered | 21 | Female | 263 | Moderna | 177 | 237 | 139 | 10 |
| 12 | Recovered | 39 | Male | 178 | Moderna | 148 | 1168 | 688 | 41 |
| 13 | Recovered | 67 | Female | 91 | Moderna | 169 | 474 | 111 | 25 |
| 14 | Recovered | 36 | Female | 369 | Pfizer | 175 | 2422 | 1723 | 308 |
| 15 | Recovered | 47 | Male | 266 | Pfizer | 162 | 171 | 103 | 10 |
| 16 | Recovered | 77 | Female | 339 | Pfizer | 191 | 633 | 280 | 50 |
| 17 | Recovered | 66 | Male | 335 | Pfizer | 191 | 3215 | 1999 | 10 |
| 18 | Recovered | 58 | Female | 361 | Pfizer | 184 | 315 | 127 | 22 |
| 19 | Recovered | 59 | Male | 388 | Pfizer | 174 | 3296 | 3102 | 423 |
| 20 | Recovered | 65 | Male | 318 | Pfizer | 182 | 431 | 247 | 42 |
| 21 | Recovered | 63 | Female | 371 | Pfizer | 175 | 615 | 508 | 30 |
| 22 | Recovered | 59 | Female | 371 | Pfizer | 177 | 2244 | 967 | 135 |
| 23 | Recovered | 54 | Male | 376 | Pfizer | 188 | 424 | 68 | 10 |
| 24 | Recovered | 40 | Female | 378 | Pfizer | 173 | 166 | 46 | 10 |
| 25 | Recovered | 49 | Male | 307 | Pfizer | 167 | 2247 | 1289 | 160 |
| 26 | Recovered | 30 | Female | 288 | Pfizer | 180 | 81 | 183 | 10 |
| 27 | Recovered | 44 | Female | 352 | Pfizer | 178 | 389 | 338 | 13 |
| 28 | Recovered | 53 | Female | 374 | Pfizer | 164 | 15 | 10 | 10 |
| 29 | Recovered | 43 | Female | 267 | Pfizer | 198 | 3075 | 1099 | 159 |
| 30 | Recovered | 52 | Male | 169 | Pfizer | 187 | 1583 | 1369 | 13 |
| 31 | Recovered | 42 | Male | 245 | Pfizer | 192 | 2688 | 812 | 11 |
| 32 | Recovered | 67 | Female | 245 | Pfizer | 184 | 4737 | 6668 | 645 |
| 33 | Recovered | 72 | Male | 235 | Pfizer | 184 | 713 | 161 | 44 |
| 34 | Recovered | 24 | Female | 171 | Pfizer | 191 | 381 | 41 | 10 |
| 35 | Recovered | 30 | Female | 96 | Pfizer | 182 | 255 | 185 | 10 |
| 36 | Recovered | 60 | Male | 40 | Pfizer | 174 | 332 | 175 | 10 |
| 37 | Recovered | 63 | Female | 40 | Pfizer | 178 | 224 | 67 | 14 |
