## Supplementary Table 4 for "mRNA-1273 and BNT162b2 mRNA vaccines have reduced neutralizing activity against the SARS-CoV-2 Omicron variant"

**Supplemental Table 4. FRNT50 results from Naïve Moderna or Pfizer vaccinated samples (1-4 weeks post 3rd dose)**

| S.No | Age | Sex | Initial vaccine | Booster dose | Interval between 2nd and 3rd dose | Days Post Dose 3 | WA1/2020 | B.1.351 | B.1.1.529 |
|---|---|---|---|---|---|---|---|---|---|
| 1 | 29 | Female | Pfizer | Moderna | 288 | 9 | 3646 | 1792 | 577 |
| 2 | 26 | Male | Pfizer | Moderna | 253 | 6 | 748 | 308 | 56 |
| 3 | 26 | Female | Pfizer | Moderna | 220 | 7 | 992 | 507 | 69 |
| 4 | 50 | Female | Pfizer | Moderna | 286 | 28 | 1335 | 605 | 175 |
| 5 | 40 | Male | Pfizer | Pfizer | 261 | 11 | 1122 | 567 | 173 |
| 6 | 28 | Male | Pfizer | Pfizer | not known | 27 | 889 | 456 | 131 |
| 7 | 33 | Female | Pfizer | Pfizer | 246 | 28 | 987 | 733 | 177 |
| 8 | 31 | Female | Pfizer | Pfizer | 260 | 28 | 2143 | 1246 | 166 |
| 9 | 29 | Female | Pfizer | Pfizer | 263 | 30 | 268 | 94 | 10 |
| 10 | 54 | Male | Pfizer | Pfizer | 264 | 29 | 1974 | 1395 | 258 |
| 11 | 26 | Female | Pfizer | Pfizer | 209 | 26 | 1141 | 667 | 127 |
| 12 | 60 | Female | Pfizer | Pfizer | 272 | 29 | 451 | 120 | 15 |
| 13 | 44 | Male | Pfizer | Pfizer | 269 | 29 | 2274 | 881 | 250 |
| 14 | 40 | Female | Pfizer | Pfizer | 268 | 32 | 3677 | 1055 | 181 |
| 15 | 53 | female | Pfizer | Pfizer | 271 | 30 | 1593 | 535 | 114 |
| 16 | 25 | Female | Pfizer | Pfizer | 248 | 31 | 949 | 358 | 45 |
| 17 | 28 | Male | Pfizer | Pfizer | 269 | 30 | 1788 | 1185 | 315 |
| 18 | 32 | Female | Pfizer | Pfizer | 273 | 30 | 1177 | 689 | 54 |
| 19 | 34 | Female | Pfizer | Pfizer | 273 | 29 | 244 | 89 | 10 |
| 20 | 27 | Female | Pfizer | Pfizer | 276 | 28 | 1522 | 603 | 76 |
| 21 | 28 | Male | Pfizer | Pfizer | 282 | 27 | 446 | 100 | 23 |
| 22 | 40 | Female | Pfizer | Pfizer | 277 | 27 | 2328 | 1358 | 360 |
| 23 | 29 | Male | Pfizer | Pfizer | 279 | 28 | 5057 | 1128 | 351 |
| 24 | 29 | Male | Pfizer | Pfizer | 264 | 27 | 736 | 230 | 73 |
| 25 | 24 | male | Pfizer | Pfizer | 261 | 31 | 3164 | 706 | 558 |
| 26 | 31 | Male | Pfizer | Pfizer | 305 | 7 | 1491 | 408 | 195 |
| 27 | 27 | Female | Pfizer | Pfizer | 273 | 25 | 649 | 175 | 62 |
| 28 | 29 | Male | Pfizer | Pfizer | 269 | 29 | 1511 | 765 | 143 |
| 29 | 38 | Female | Pfizer | Pfizer | 291 | 29 | 857 | 411 | 32 |
| 30 | 25 | Male | Pfizer | Pfizer | 209 | 30 | 629 | 199 | 43 |
| 31 | 35 | Female | Pfizer | Pfizer | 253 | 33 | 1075 | 264 | 10 |
| 32 | 65 | Female | Pfizer | Pfizer | 269 | 30 | 1866 | 484 | 116 |
| 33 | 30 | Female | Pfizer | Pfizer | 209 | 30 | 1014 | 395 | 46 |
| 34 | 20 | Male | Pfizer | Pfizer | 233 | 27 | 1095 | 378 | 44 |
| 35 | 26 | Female | Pfizer | Pfizer | 286 | 28 | 5206 | 1260 | 160 |
| 36 | 51 | Male | Moderna | Moderna | 251 | 29 | 1561 | 975 | 197 |
| 37 | 26 | Female | Moderna | Moderna | 210 | 7 | 1042 | 289 | 53 |
| 38 | 26 | Female | Moderna | Moderna | 256 | 28 | 2808 | 1317 | 264 |
| 39 | 33 | Female | Moderna | moderna | 257 | 8 | 717 | 520 | 71 |
| 40 | 27 | Female | Moderna | Moderna | 278 | 7 | 1317 | 432 | 74 |
| 41 | 68 | Female | Moderna | Moderna | 197 | 7 | 1629 | 1233 | 110 |
| 42 | 67 | Female | Moderna | Moderna | 274 | 7 | 1504 | 639 | 56 |
| 43 | 33 | Male | Moderna | Moderna | 292 | 7 | 699 | 299 | 54 |
| 44 | 37 | Female | Moderna | Moderna | 269 | 28 | 2351 | 1112 | 145 |
| 45 | 30 | Female | Moderna | Moderna | 247 | 28 | 2890 | 1923 | 185 |
| 46 | 32 | Male | Moderna | Moderna | 222 | 35 | 1556 | 814 | 69 |
| 47 | 79 | Male | Moderna | Moderna 100mcg | 256 | 7 | 3137 | 1495 | 235 |
| 48 | 72 | Male | Moderna | Moderna 100mcg | 223 | 30 | 1250 | 486 | 71 |
| 49 | 27 | Male | Moderna | Pfizer | 281 | 6 | 165 | 55 | 10 |
| 50 | 27 | Female | Moderna | Pfizer | 277 | 9 | 3346 | 2206 | 375 |
| 51 | 28 | Female | Moderna | Pfizer | 291 | 7 | 1028 | 258 | 57 |
| 52 | 38 | Female | Moderna | Pfizer | 225 | 32 | 1654 | 1230 | 148 |
